# HBM-CITEseq: a uniform CITE-seq processing pipeline for the HuBMAP Consortium

**DOI:** 10.1101/2022.12.19.521058

**Authors:** Xinyue Lu, Matthew Ruffalo

## Abstract

As part of the HuBMAP Consortium we have been developing methods for uniformly processing and indexing multiple single-cell datasets, which enable efficient integration of data from different platforms. HuBMAP computational pipelines have so far focused on unimodal data types such as single-cell/nucleus RNA sequencing and single nucleus ATAC-seq. Here we present HBM-CITEseq, a processing pipeline for Cellular Indexing of Transcriptomes and Epitopes by Sequencing (CITE-seq) datasets, the first multimodal sequencing processing pipeline for the HuBMAP Consortium, with transcriptomic outputs from HBM-CITEseq matching those of the HuBMAP RNA-seq pipeline.

HBM-CITEseq is a CWL workflow wrapping command-line tools encapsulated in Docker images. It is freely available on GitHub at https://github.com/hubmapconsortium/citeseq-pipeline.

## Introduction

Single cell RNA-sequencing data (scRNA-seq) can provide useful information on highly heterogeneous cell populations across multiple tissues, disease states, time points, etc. However, different projects utilize different technologies and data processing methods, which raises significant practical challenges in integrative data analysis. To avoid these difficulties, the Human BioMolecular Atlas Program [3] (HuBMAP Consortium) performs uniform processing of single-cell, single-nucleus, and spatially resolved RNA-seq data, using a standardized computational pipeline (https://github.com/hubmapconsortium/salmon-rnaseq). Supported assays include 10X Genomics Chromium v2 and v3, SNARE-seq, sci-RNA-seq, Slide-seq, and more, with new assays added continuously to match data types provided by HuBMAP tissue mapping centers and other data providers.

Cellular Indexing of Transcriptomes and Epitopes by Sequencing (CITE-seq) [6] is a single-cell sequencing approach that can make up the limitation for current scRNA-seq sequencing technologies that mRNA expression level can not accurately reflect protein abundance due to post-transcriptional modifications. CITE-seq utilizes DNA-barcoded antibodies to corporate both transcriptome-wide measurements for single cells and protein expression at cell-surface level, which enables downstream multimodal analysis. scRNA-seq library preparation protocols used for capturing DNA-barcoded antibodies include 10x Genomics, Drop-seq, ddSeq, etc.

Here, we introduce an additional CITE-seq pipeline HBM-CITEseq which takes hash-tag barcoding antibody-oligos (HTO) data as an optional input and integrates RNA expression and CITE-seq antibody-oligos (ADT) data for multi-omics analysis. In detail, HBM-CITEseq utilizes the HuBMAP scRNA-seq pipeline as a submodule to quantify sc-RNA expression data. It can also handle the indexing and quantification of ADT and HTO data. Multi-omics integration is also performed on the multimodal data.

## Implementation

HBM-CITEseq is a CITE-seq processing pipeline built on Salmon [5], Scanpy [8] and Muon [2]. It is an extension of the HuBMAP scRNA-seq pipeline (https://github.com/hubmapconsortium/salmon-rnaseq) which can uniformly process and analyze CITE-seq data.

In addition to the input of the HuBMAP scRNA-seq pipeline, HBM-CITEseq requires the FASTQ directory containing raw data and feature barcode information for ADT data and HTO data (optional). If the feature barcoding protocol utilizes TotalSeq B or C, a barcode transformation mapping file is also required to match the cellular barcodes of RNA with feature barcode library.

HBM-CITEseq has separate quantification steps for RNA, ADT and HTO. For RNA quantification, HBM-CITEseq utilizes the HuBMAP scRNA-seq pipeline as a submodule. For ADT and HTO quantification, it includes the following steps: (1) Indexing the feature barcodes to quantify the antibodies and the sample multiplexing based cell-hashing. (2) Run alevin to quantify the gene abundances based on the relevant 3-prime sequencing technology with the previously generated index. `--naiveEqclass` flag is extended for the UMI deduplication of the HTO data. (3) Convert the unfiltered count matrix from Alevin to H5AD. Finally, a Muon object is constructed using the RNA data, ADT and HTO (optional) data is added as a separate assay to the object for the common cells. The RNA and ADT modalities in Muon format are used for downstream analysis. To analyze each individual modality, HBM-CITEseqperforms: (1) Normalization and feature selection. (2) PCA and neighborhood graph computation. (3) Non-linear dimensionality reduction and clustering. To investigate the multi-omics integration, HBM-CITEseq performs: (1) Multi-omics factor analysis(MOFA) [1], which generates an interpretable latent space jointly on both modalities. (2) Compute the cell neighborhood graph using MOFA factors learned on both modalities. [4]

Together, HBM-CITEseq facilitates the uniform quantification and interpretation of CITE-seq data. The output multimodal object can be further used to perform cell type annotation and MOFA model interpretation.

## Results

Testing on GSE128639 [7] shows that HBM-CITEseq can generate comparable quantification and downstream analysis results compared with the public processed data, with sc/snRNA-seq results directly compatible with datasets available on the HuBMAP portal (same reference transcriptome, quantification method, genome annotations, etc.). Figure 1 (B), (C) 1 show the Leiden clustering results on RNA and ADT modalities for public processed data and our processed data. For more details, please see Supplementary Material.

**Figure 1.**
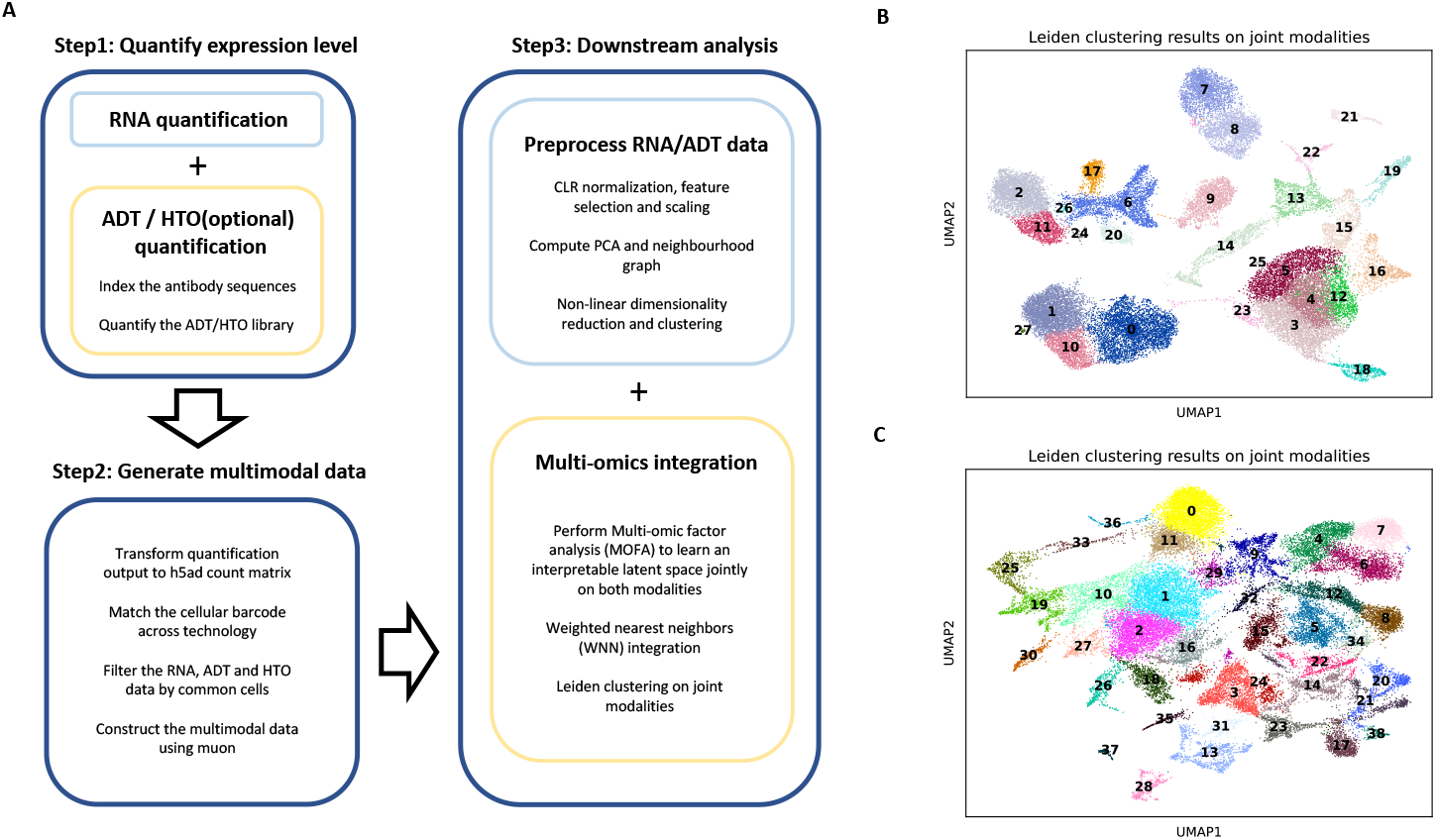
(A) Overview of HBM-CITEseq for CITE-seq processing. (B) Joint embedding results on RNA and ADT modalites for public processed data on GEO. (C) Joint embedding results on RNA and ADT modalites for HBM-CITEseq processed data

## Conclusion

Being an extension of HuBMAP scRNA-seq pipeline, HBM-CITEseq enables a uniform way to index, process and visualize CITE-seq sequencing data for the HuBMAP Consortium. It allows for integration of gene expression and surface protein abundance, which can be further used for multimodal downstream analysis.

## Acknowledgements

The authors would like to acknowledge computational support provided by the Pittsburgh Supercomputing Center as part of the Human BioMolecular Atlas Program (HuBMAP).

