## Supplementary File for "HBM-CITEseq: a uniform CITE-seq processing pipeline for the HuBMAP Consortium"

### HuBMAP CITE-seq pipeline supplement

Xinyue Lu [1,*] and Matthew Ruffalo [1,*]

September 2022

#### 1 Comparison of embedding results between public processed data and our processed data on GSE128639

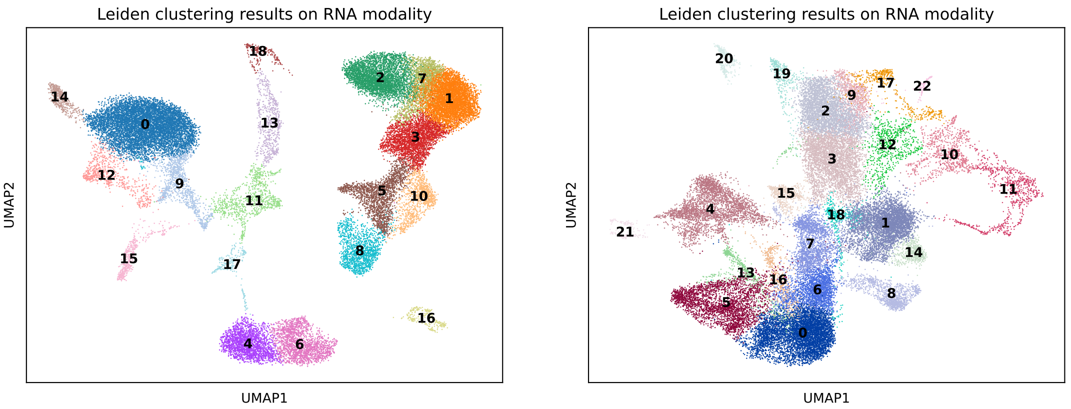


(a) Public processed data

(b) Our processed data

Figure 1: Comparison of leiden clustering results on RNA modality

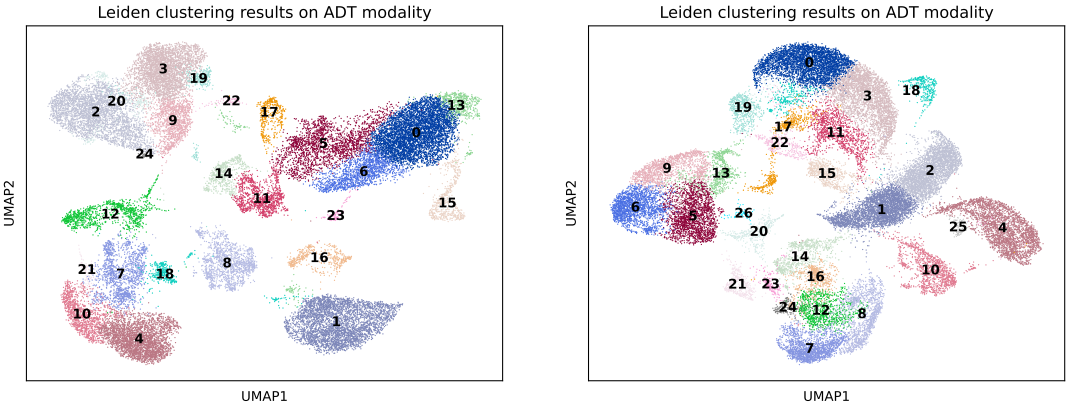


(a) Public processed data

(b) Our processed data

Figure 2: Comparison of leiden clustering results on ADT modality

### 2 Comparison of expression level between public processed data and our processed data on GSE128639

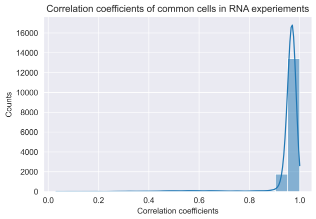


(a) RNA modality

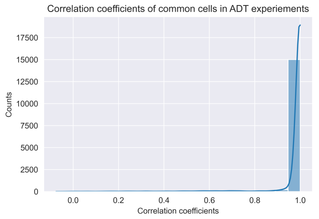


(b) ADT modality

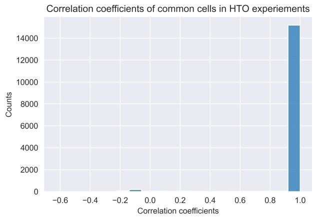


(c) HTO modality

Figure 3: Distribution of correlation coefficient in common cells of public processed data and our processed data. The correlation coefficients are high in all modalities.

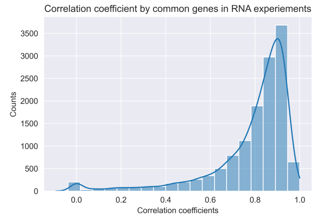


(a) RNA modality

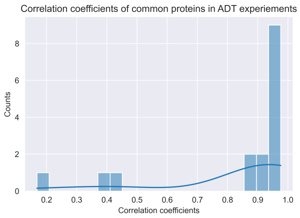


(b) ADT modality

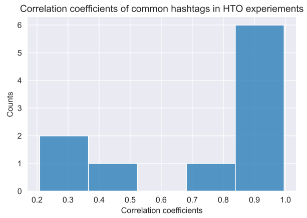


(c) HTO modality

Figure 4: Distribution of correlation coefficient in common genes/proteins/hashtags of public processed data and our processed data. The correlation coefficients are high in all modalities.

#### 3 Comparison of barcode occurrence of identified cells in raw FASTQ file between public processed data and our processed data on GSE128639

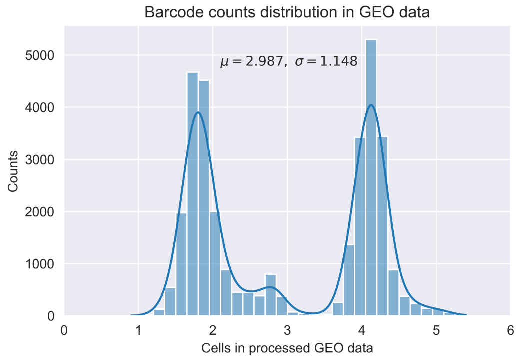


(a) Public processed data

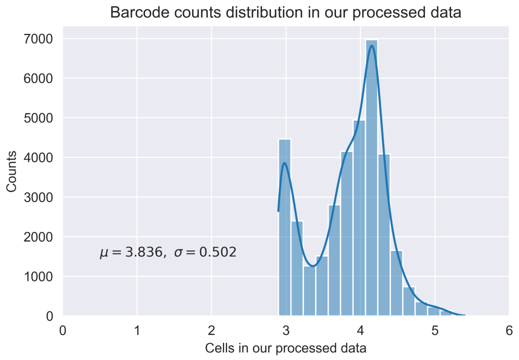


(b) Our processed data

Figure 5: Barcode occurrence of identified cells in raw FASTQ file before error correction.
